# CTPC, a combined transcriptome dataset of prostate cancer cell lines

**DOI:** 10.1101/2022.07.06.499012

**Authors:** Siyuan Cheng, Xiuping Yu

**Affiliations:** Department of Biochemistry & Molecular Biology, LSU Health Shreveport; Department of Urology, LSU Health Shreveport

## Abstract

Cell lines are the most used model system in cancer research. The transcriptomic data of established prostate cancer (PCa) cell lines help researchers to explore the differential gene expression across the different PCa cell lines and develop hypothesis. Through large scale datamining, we established a curated <u>C</u>ombined <u>T</u>ranscriptome dataset of <u>P</u>Ca <u>C</u>ell lines (CTPC) which contains the transcriptomic data of 1840 samples of seven commonly used PCa cell lines including LNCaP, C4-2, VCaP, 22Rv1, PC3, DU145, and NCI-H660. The CTPC dataset provides an opportunity for researchers to not only compare gene expression across different PCa cell lines but also retrieve the experiment information and associate the differential gene expression with meta data such as gene manipulation and drug treatment information. Additionally, based on the CTPC dataset, we built a platform for users to visualize the data (https://pcatools.shinyapps.io/CTPC_V2/). It is our hope that the combined CTPC dataset and the user-friendly platform are of great service to the PCa research community.

## Background & Summary

Prostate cancer (PCa) is one of the most common cancers in American men and the second leading cause of cancer-related deaths. Cell line models are commonly used in PCa research. They are cost efficient and relatively easy to maintain and genetically manipulate. The transcriptomic data of PCa cell lines can provide invaluable molecular information for researchers to choose suitable cell lines for their research projects or develop testable hypotheses, as different cell lines can reflect the disease at different stage.^1^ Two pan-cancer studies, Cancer Cell Line Encyclopedia (CCLE) and NCI-60^2,3^, have generated transcriptomic data for many cancer cell lines including several PCa cell lines. However, both datasets lack replicates, which weakens the data’s reliability. Gene Expression Omnibus (GEO) database has collected a large amount of transcriptomic data derived from PCa cell lines, but these data were generated by different research groups, following different experimental protocols and using different sequencing platforms. Basic bioinformatic skills are needed to retrieve gene expression data from these studies, and more advanced skills are needed to compare data across different studies. We established a curated Combined Transcriptome dataset of PCa Cell lines (CTPC) which contains 1840 samples covering seven commonly used PCa cell lines (LNCaP and its derivative C4-2, VCaP, 22RV1, PC3, DU145 and NCI-H660). These cells were cultured under various conditions or had undergone certain genetic alterations. Additionally, we developed a user-friendly platform to visualize the normalized gene expression data as well as the annotated meta data of the samples collected in CTPC (https://pcatools.shinyapps.io/CTPC_V2/). This allows scientists to examine the expression of the genes of their interest across different PCa cell lines and correlate gene expression levels with treatment conditions.

## Methods

The datamining was performed using R/ Rstudio^4,5^ with customized codes. The raw sequencing data were extracted from ARCHS^4^, which stores a collection of gene expression count matrixs adapted from GEO^6^, using key words “VCaP”, “LNCaP”, “C42”, “22RV1”, “PC3” and “DU145”. The transcriptomic data of NCI-H660, the solely available neuroendocrine prostate cancer cell line, were manually extracted from the GEO database using keyword “H660”as ARCHS4 database collected few samples of this cell line. A total of 14 H660 samples from 6 GEO studies were found. The RNA-seq data generated by CCLE and Dr. Korkola’s group were also included because both contain H660 samples^2,7^. Among all the 1840 samples, 15 samples from 3 studies were in “FPKM” format and the rest 1825 samples were in “Counts” format. The “Count” data were first converted into FPKM format using the “fpkm” function in “DEseq2” package. The mRNA size vector was retrieved from ENSEMBL database^8^ and the 1840 “FPKM” samples were merged, log2 transformed and normalized. The meta data including “cell line”, “treatment” and “Resource” were manually extracted from the original publications or GEO website based on the samples’ “GEO sample number” and “GEO series number”. The mislabelled samples, falsely selected samples, and those that were sequenced by method other than the commonly used bulk RNA-seq (for example GRO-seq) were removed from the dataset. The normalized FPKM expression data and meta data of the 1840 samples were combined into a single dataset which is designated as “CTPC dataset”. Additionally, to help researchers visualize the gene expression data without R coding, we built a user-friendly online platform using the “Shiny” package^9^ (https://pcatools.shinyapps.io/CTPC_V2/). The “ggplot” and “plotly” packages were used to generate the interactive violin plots^10,11^.

## Data Records

A total of 175 GEO studies were included in the CTPC dataset. Seven PCa cell lines with 344 different treatments were used in these studies. The sample numbers for each cell line are 181 for 22RV1, 147 for C4-2, 83 for DU145, 906 for LNCaP, 279 for PC3, 228 for VCaP and 16 for H660. “WT” represents the non-treated cells and “Control” represents the control samples including vehicle control, scramble and nontargeting RNA controls. Both meta data and the GEO accession number could be retrieved by clicking the data point on the interactive plots. This provides an opportunity for researchers to obtain more information on their study of interest.

## Technical Validation

For technical validation, we generated boxplots using the log2 FPKM values. The samples were presented individually (Figure 1A) or grouped by experimental batches (Figure 1B) or cell lines (Figure 1C). As shown in these figures, both the distribution (1^st^ quarter to 3^nd^ quarter) and median values appear similar across all the samples in CTPC, indicating a good normalization of the transcriptomic data. Also, we conducted dimensional reduction on the CTPC dataset. The result was visualized using UMAP. As shown in Figure 1D, samples of the same PCa cell line tend to co-locate and samples of different cell lines are clearly separated except LNCaP and C4-2 samples that cluster together. This is not surprising since C4-2 line is a LNCaP-derivative^12^. Additionally, we highlighted the expression of two genes (MYCN and CPT1A) on the boxplots. As shown in Figure 1E, the LNCaP and 22RV1 samples that ectopically express MYCN exhibit high levels of MYCN mRNA. Similarly, the C4-2/CPT1A overexpression samples displayed high levels of CPT1A mRNA while the C4-2/CPT1A knock-down samples (shCPT1A) displayed low expression (Figure 1F).

**Figure 1.**
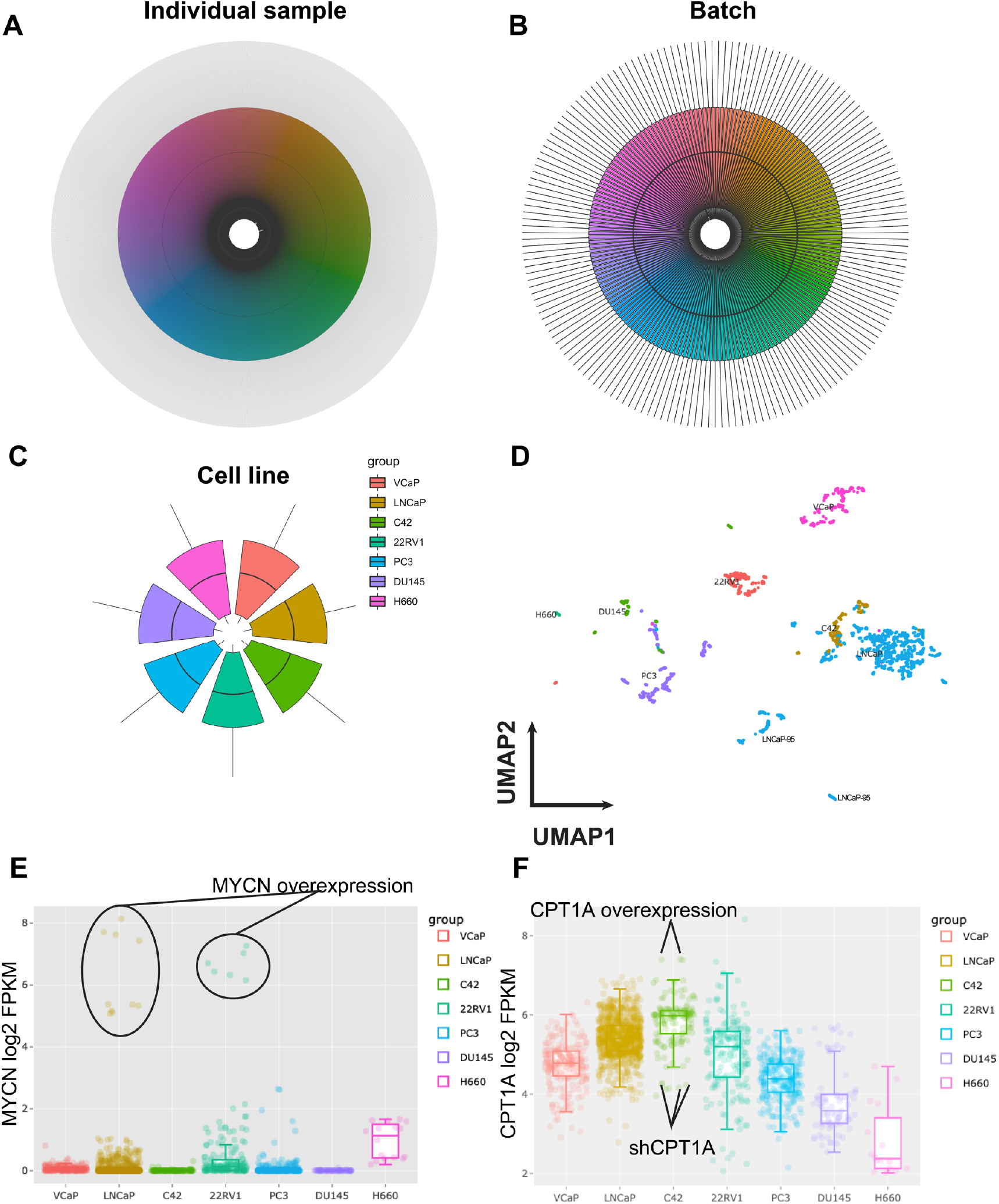
quality validations of the CTPC dataset. (A) Circular boxplot of mRNA FPKM value of individual samples in the CTPC dataset. The X axis (circumference) represents the 1840 samples (in different colors) and Y axis (radius) is the log2 FPKM value. (B) Circular boxplot of the 1840 CTPC samples grouped by batches. (C) Circular boxplot of the samples grouped by cell lines. (D) UMAP clusters of the CTPC dataset. The seven PCa cell lines were labelled with different colors. The samples from the same cell line tend to co-locate. (E and F) Boxplots generated using the interactive online platform. The samples that ectopically express MYCN, CPT1A or-shCPT1A were highlighted. The mRNA expression of these genes matches the genetic manipulation of the cells.

## Usage Notes

The CTPC dataset is the first comprehensive RNASeq dataset that focuses on PCa cell lines. This provides access to gene expression data across PCa cell lines with sufficient sample numbers. Additionally, the online platform enables researchers to conduct integrative analyses of gene expression in each PCa cell line, compare gene expression levels across different cell lines, and correlate the altered gene expression with treatment conditions. The GEO accession number is also readily available for each sample. We believe the CTPC dataset and associated platform provide a great service to the PCa research community and will enable the development of many exciting and impactful research projects.

## Code Availability

All custom R codes are available upon request.

## Author contributions

S. Cheng: Conceptualization, Software, data collection, data visualization, and manuscript writing

X. Yu: project supervision, funding acquisition, and manuscript revision.

## References

1 Cheng, S. & Yu, X. The spectrum of neuroendocrine differentiation in prostate cancer. Prostate Cancer and Prostatic Diseases 24, 1214–1215 (2021).

2 Ghandi, M. et al. Next-generation characterization of the Cancer Cell Line Encyclopedia. Nature 569, 503 (2019).

3 Reinhold, W. C. et al. CellMiner: a web-based suite of genomic and pharmacologic tools to explore transcript and drug patterns in the NCI-60 cell line set. Cancer research 72, 3499–3511 (2012).

4 Team, R. C. (2018).

5 Team, R. (2019).

6 Lachmann, A. et al. Massive mining of publicly available RNA-seq data from human and mouse. Nature communications 9, 1–10 (2018).

7 Smith, R. et al. Enzalutamide response in a panel of prostate cancer cell lines reveals a role for glucocorticoid receptor in enzalutamide resistant disease. Scientific reports 10, 1–13 (2020).

8 Love, M. I., Huber, W. & Anders, S. Moderated estimation of fold change and dispersion for RNA-seq data with DESeq2. Genome biology 15, 1–21 (2014).

9 Chang, W., Cheng, J., Allaire, J., Xie, Y. & McPherson, J. Shiny: web application framework for R. R package version 1, 2017 (2017).

10 Wickham, H. ggplot2: elegant graphics for data analysis. (Springer, 2016).

11 Sievert, C. Interactive web-based data visualization with R, plotly, and shiny. (CRC Press, 2020).

12 Liu, A. Y. et al. Lineage relationship between LNCaP and LNCaP - derived prostate cancer cell lines. The Prostate 60, 98–108 (2004).

